# Personality traits change after an opportunity to mate

**DOI:** 10.1101/2020.03.02.973693

**Authors:** Chloé Monestier, Alison M. Bell

**Author notes:** Corresponding author. E-mail address (C. Monestier).

## Abstract

There is growing evidence that personality traits can change throughout the life course in humans and nonhuman animals. However, the proximate and ultimate causes of personality trait change are largely unknown, especially in adults. In a controlled, longitudinal experiment, we tested whether a key life event for adults – mating – can cause personality traits to change in female threespine sticklebacks. We confirmed that there are consistent individual differences in activity, sociability and risk taking, and then compared these personality traits among three groups of females: 1) control females; 2) females that physically mated; 3) females that socially experienced courtship but did not mate. Both the physical experience of mating and the social experience of courtship caused females to become less willing to take risks and less social. To understand the proximate mechanisms underlying these changes, we measured levels of excreted steroids. Both the physical experience of mating and the social experience of courtship caused levels of dihydroxyprogesterone (17α,20β-P) to increase, and females with higher 17α,20β-P were less willing to take risks and less social. These results provide experimental evidence that personality traits and their underlying neuroendocrine correlates are influenced by formative social and life-history experiences well into adulthood.

## INTRODUCTION

An outstanding question in both the human and animal personality literature concerns the stability of individual differences throughout the life course. By definition, personality traits are relatively consistent across situations and/or over time [1,2], but they are not immutable, even in adults. For example, studies in humans have found significant mean-level change in all trait domains at some point during the life course [3,4]; people become more conscientious, warmer and calmer after the age of 30 [3]. Personality traits might change over ontogeny due to intrinsic maturation [4] and/or because juveniles and adults experience different environments, including different social roles that might favor different behavioral strategies. Alternatively, or in addition, personality traits might change following a particular experience because the experience exposes the animal to different selection pressures, and/or represents an important life history decision that influences residual reproductive value. For example, important events such as metamorphosis [5,6], migration, dispersal, sexual maturation [7,8], reproduction, parenting, etc. can expose individuals to different environments and selective pressures, thereby driving changes in personality traits.

Our understanding of the adaptive significance of personality trait change throughout the life course is limited because the causes of personality trait change are challenging to study. For one, self-selection can be a problem because some behavioral types of individuals might be more likely to experience a particular life-history event than others [9,4]. Second, even if all individuals experience the particular event, they might do so at different times or at different ages; therefore differences between those that did versus did not experience the event could reflect the confounding effects of time, age, season or maturation [10,11]. Moreover, it can be difficult to pinpoint the exact causes of personality trait change following a key life-history event because life-history events often comprise a series of sub-events leading up to them, making it difficult to isolate the effects of becoming a parent, for example, from the effects of courtship, mating, and reproduction, all of which need to occur before parenting can begin.

To circumvent these problems, here, we directly measured the impact of a specific life event - mating - on personality trait development in adult female threespine sticklebacks (*Gasterosteus aculeatus*) in a controlled, randomized and longitudinal experiment. Among life-history events, the experience of mating and reproducing for the first time is likely to be one of the most important life events for any organism. For example, courtship and/or sexual experience often influences subsequent female mate preference [12,13]; females generally become more selective with mating experience [13]. Therefore, we might expect them to become more cautious, i.e. less bold and aggressive, after mating. On the other hand, we might expect females to become more bold after mating because they have less to lose, i.e. their residual reproductive value is lower. In general, recent theory on the adaptive evolution of personality traits predicts that a female’s recent mating and courtship experience will influence her willingness to take risks in the future [14,15].

Here, we examine the influence of mating for the first time on personality trait development in female sticklebacks. Sticklebacks are famous for their natural intraspecific variation in behavior [16,17], and a previous study found that the experience of reproduction and parenting influenced the development of risk taking behavior in male sticklebacks [18]. In contrast to males, female sticklebacks do not provide parental care. Instead, female sticklebacks become gravid and lay eggs in males’ nests, where they are fertilized externally [17,19]. Sticklebacks from most freshwater populations die at the end of their first and only breeding season [17,19].

We repeatedly measured personality traits on individual females and compared three groups of females: 1) females that did not have an opportunity to mate (control); 2) females that physically mated; 3) females that had an opportunity to mate and experienced courtship socially but did not mate (courtship control). Personality traits were repeatedly measured in all groups of females both before and after females had an opportunity to mate. This experimental design allowed us to determine if the physical act of mating and reproducing is required to cause personality traits to change, or if the social experience of courtship is sufficient. In order to track the potential proximate mechanisms underlying changes in personality traits as a function of mating, we used a noninvasive method to measure levels of steroids excreted in the water both before and after a mating opportunity [20]. Given the dramatic neuroendocrine changes associated with reproduction [21,22], we predicted that females that mated would experience greater steroid fluctuations compared to females that had not mated, and that those hormonal fluctuations would be related to behavior. We focused on cortisol because of its hypothesized link to personality variation [23] and on steroids involved in reproduction because it was the life history event of interest.

## METHODS

Threespine sticklebacks were collected from Putah Creek, California, U.S.A.. Neither males nor females showed signs of reproductive maturity therefore we assume that females were virgins at the time of the experiment. Females were housed in all-female groups in ‘home’ tanks (35.5 L × 33 W × 25 H cm, 10 fish/tank) with a gravel bottom, plastic plants, and an opaque shelter and stayed in these home tanks for the duration of the experiment, except when individuals were transferred to separate ‘observation’ tanks (60.75 L × 30 W × 20 H cm, set up the exact same as the home tanks) for behavior trials. Fish were maintained at 20°C on a summer photoperiod (16: 8 h light: dark cycle). The fish were daily fed a mixed diet consisting of frozen bloodworm, brine shrimp, and Mysis shrimp ad lib each day.

### Experimental design

Individual females were randomly assigned to either the control group or to have a mating opportunity. Females given a mating opportunity were paired with a control female (matched for size) who was always measured at the same time as her partner. This design allowed us to control for variation among females in time to become gravid and to reproduce, as well as for self-selection, i.e. if some behavioral types of females are less choosy or more attractive than others. Females in the control group (n = 37) were not exposed to a male or given an opportunity to mate, but like females in the other conditions, they were socially housed, therefore differences between control females and females given a mating opportunity do not reflected differences in the opportunity for social interactions per se. Moreover, many females (n = 12) in the control group also became gravid over the course of the experiment and released their eggs in their home tank. We tested for the effects of releasing eggs on behavior by comparing the behavior of control females before vs after releasing eggs, and there was no effect (activity: F_(1,220)_ = 2.74, p-val = 0.10; social behavior: F_(1,220)_ = 1.57, p-val = 0.19; risk-taking behavior: F_(1,220)_ = 0.05, p-val = 0.82; 17α,20β-P: F_(1,12)_ = 0.001, p-val = 0.99). Therefore, behavioral differences between females in the control group and females given a mating opportunity do not reflect differences in gravidity, or the effects of releasing eggs. Some of the females given a mating opportunity (n = 23) mated, while others (n = 22) did not and served as “courtship controls”, because like the mated females, they had the social experience of being courted, but unlike the mated females they did not physically mate. We did not detect any latent differences between females that mated and courtship control females that would lead to nonrandom representation of behavioral types between the mated and courtship control groups, e.g. mated and courtship control females did not differ in size (t_43.95_ = 0.16, P = 0.87) or behavior (see Results). Both courtship control and mated females had multiple opportunities to mate (mated females: range = 1 – 8 opportunities, mean ± SE = 2.9 ± 0.29, courtship control females: range = 1 – 7, mean ± SE = 3.4 ± 0.31).

The behavior of females was measured in the behavioral assays on six occasions, thrice in the “Before” trials and thrice in the “After” trials, with 24 h between trials. Females given a mating opportunity were placed in a tank with a male who had built a nest (60.75 L × 30 W × 20 H cm). Females started the After behavioral assays 24 hours after the mated female reproduced or after the courtship control female released her eggs in her home tank (presumably to avoid the costs of becoming egg bound [24]. All females were measured for length and weight on the last day of the After trials.

### Behavioral assays

#### Activity

The focal fish was placed in a shelter at one end of the tank. After one minute, the observer gently removed the cork of the shelter, and recorded the number of areas (four 15 cm squares) crossed for three minutes.

#### Social behavior

30 minutes after the activity assay, three females from the same population were placed into a flask at the opposite side of the refuge and we recorded the number of times the focal fish contacted the flask for 5 minutes.

#### Risk taking behavior

30 minutes after the social behavior assay, a model great egret (*Casmerodius albus*) head was placed over the observation tank. Then, we added live bloodworms directly under the egret. When the focal fish approached the worms within one body length, we released the egret twice in quick succession, and then fixed the egret so that it remained above the water. Following this simulated attack, we recorded time foraging under risk for five minutes.

### Measuring excreted steroids

After the third trial of both the Before and After trials, we placed the focal fish in a 500 ml long-necked glass flask filled with 100 ml of water. We then placed the flask in a covered bucket (to minimize stress) for 30 minutes. Then, we transferred 50 ml of the water into a 50 ml sterile polypropylene conical tube.

Steroids were extracted from the water samples by pulling water through C18 Sep-Pak cartridges (Waters Ltd.) that had been primed with 5 mL methanol followed by 5 mL distilled water. After the sample had dripped through at a rate of approximately 2 mL/min, the cartridge was washed with 5 mL of distilled water, and the steroids were eluted from the columns into 13*100 mm borosilicate vials via 5 ml of diethyl ether. The ether was dried by evaporation overnight. The dried hormones were then frozen at −80°C.

Samples were analyzed via mass spectrometry with the 5500 QTRAP LC/MS/MS system (AB Sciex, Foster City, CA). In order to control for differences in body size, hormone release rates were calculated as the amount of released hormone per gram of body weight per hour (ng/g/hr). We focus here on 17α,20β-P (17α,20β-dihydroxy-4-pregnen-3-one, hereafter referred to as 17α,20β-P, n = 19 individuals with one measure Before and one measure After).

### Statistical analyses

#### Repeatability of behavior and hormones

To confirm that there were consistent individual differences in behavior, we estimated repeatability during the Before trials. Repeatability for the three treatment groups was estimated separately to confirm that all groups showed similar patterns prior to the mating opportunity. To test whether mating influences rank-order stability, we estimated the repeatability of behavior and hormone titres across the six Before and After trials for the treatment groups separately.

Repeatability was estimated as the ratio of between-individual variance to total variance with linear mixed-effects models (with individual identity as a random factor) in R v.3.4.4 (http://www.r-project.org) [25].

#### How do personality traits and hormones change following mating?

To detect mean-level personality trait change after mating, we used linear mixed-effects models (LMMs) with the lmer function in the R package lme4 v.1.1-17 [26]. Models included the following fixed effects: treatment (three levels: control, courtship control, and mated), Before/After (two levels: before vs after mating), trial nested within Before/After, the number of days between the first and sixth trial (days in the experiment), the number of mating attempts, the interaction between treatment x Before/After, and individual as a random effect.

We used LMMs to investigate changes in hormonal release rate according to the same explanatory variables cited above. However, we split the all data set by the period (Before/After) as we did not have enough statistical power to test for the interaction between treatment and period. We used LM to investigate a potential link between hormones and personality traits (average behavior across the three before or after trials).

More details about the methods (behaviors recorded, hormones measured and their relevance, detailed protocols and technics, and statistics) are in electronic supplementary material.

## RESULTS

### Repeatability of behavior

During the Before trials, individual differences in behavior were repeatable in all treatment groups (electronic supplementary material, Table S1), confirming these behaviors can be considered personality traits. Among-individual variation was consistently higher than within-individual variation.

Individual differences in behavior were also repeatable between the Before and After trials (Table 1) indicating some element of stability of individual behavioral types throughout the experiment. However, the repeatability of social behavior and risk taking behavior before and after the mating opportunity was significantly lower in both the mated and courtship control treatment groups compared to the control group (Table 1). This pattern appears to reflect greater among-individual variation in the control group, and higher within-individual variation in the mated and courtship control groups (Table 1). Greater within-individual variation in the mated and courtship control groups is visually evident in the behavioral reaction norms (electronic supplementary material, Fig. S1), especially between trials 3 and 4, i.e. the interval during which females in these groups experienced courtship and/or mating. The repeatability of activity did not differ among the three treatment groups (Table 1).

**Table 1.** Repeatability (R) and variance components (among- and within-individual) of behavioral traits across the before and after trials.

| Treatment groups | Activity:<br>number of areas crossed |  |  | Social behavior:<br>number of contacts |  |  | Risk taking behavior:<br>time foraging under risk |  |  |
| --- | --- | --- | --- | --- | --- | --- | --- | --- | --- |
|  | Among | Within | R | Among | Within | R | Among | Within | R |
| Control (n=37) | 54.6 | 31.17 | 0.64 (0.48, 0.74) | 15.46 | 10.58 | 0.59 (0.43, 0.71) | 4332 | 1051 | 0.81 (0.71, 0.87) |
| Mated (n=22) | 34.81 | 44.89 | 0.44 (0.22, 0.59) | 7.11 | 14.27 | 0.33 (0.12, 0.51) | 3158 | 3935 | 0.45 (0.22, 0.61) |
| Courtship control (n=23) | 32.47 | 27.35 | 0.53 (0.32, 0.69) | 6.54 | 14.78 | 0.31 (0.12, 0.49) | 2775 | 5059 | 0.35 (0.14, 0.53) |

### Effects of a mating opportunity on behavior

We did not detect any differences among the treatment groups in activity during the Before (F_(2,243)_ = 2.90, p-val = 0.10) or After trials (F_(2,243)_ = 0.48, p-val = 0.62), and no difference in activity between the Before and After trials (F_(2,489)_ = 2.29, p-val = 0.13) (Fig. 1A).

**Figure 1.**
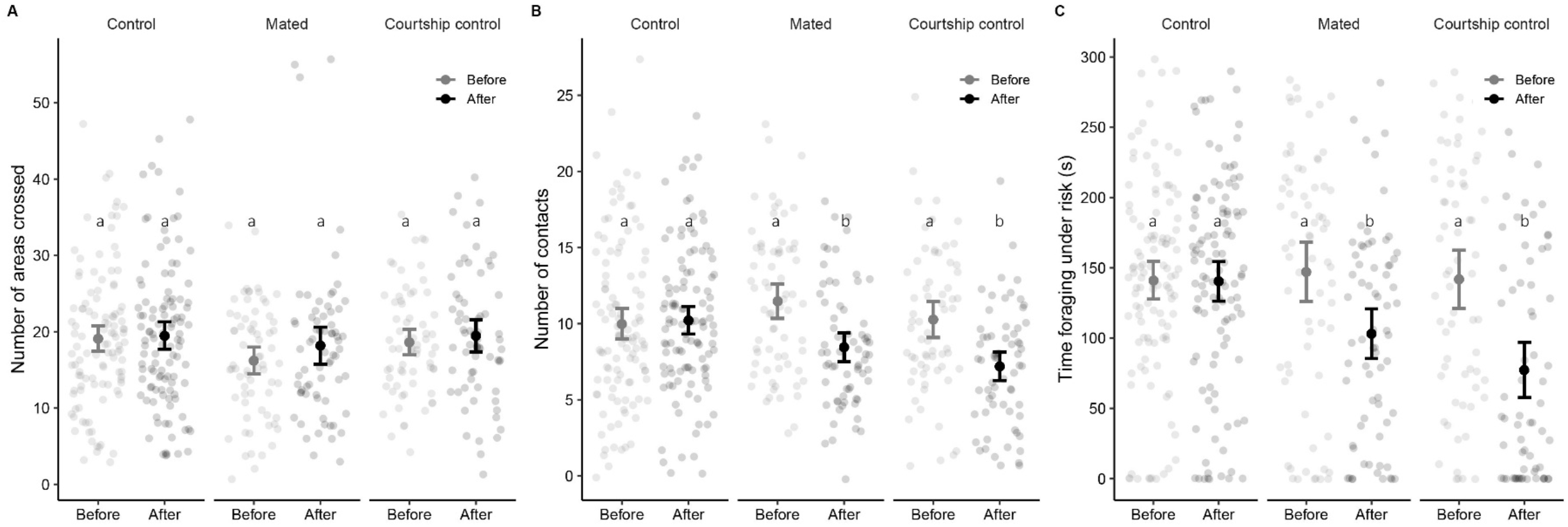
Mean-level change in behavior as a function of courtship and mating. Bars represent 95% confidence intervals. Different letters within each figure indicate means that are significantly different (p-val < 0.05).

Females in both the mated and courtship control groups were less social (fewer contacts with conspecifics) after they were given a mating opportunity (t_131.97_ = −4.09, P < 0.001; t_123.21_ = −4.07, P < 0.001, respectively), but the social behavior of females in the control group did not change across the experiment (t_217.44_ = 0.34, P = 0.73, Fig. 1B). The mated and courtship control females did not differ in social behavior (t_263.19_ = −1.54, P = 0.12). Females in the three treatment groups did not differ in social behavior during the Before trials (F_(2,243)_ = 1.93, p-val = 0.15).

Time foraging under risk was lower in both the mated and courtship control groups after they were given a mating opportunity (t_131.81_ = −3.19, P < 0.001; t129.63 = −4.52, P < 0.001, respectively), but the time that control females spent foraging under risk did not vary across the experiment (t_219.67_ = −0.09, P = 0.93, Fig. 1C). The mated and courtship control groups did not differ (t_265.57_ = −1.4, P = 0.13). Females in the three treatment groups did not differ in risk taking behavior during the Before trials (F_(2,243)_ = 0.13, p-val = 0.88).

Non-significant results about the other explanatory variables are in the electronic supplementary material.

### 17α,20β-P was higher after a mating opportunity and was negatively correlated with social behavior and risk taking behavior

The experience of courtship caused 17α,20β-P to increase: 17α,20β-P did not differ among the three treatment groups during the Before trials (F_(2,16)_ = 1.69, p-val = 0.22), but 17α,20β-P was significantly higher during the After trials in both the mated and courtship control groups compared to the control group (F_(2,16)_ = 3.49, p-val = 0.04, electronic supplementary material Fig. S2).

We did not detect a relationship between behavior and 17α,20β-P during the Before trials (number of contacts: F_(1,17)_ = 0.42, p-val = 0.52; willingness to forage under risk: F_(1,17)_ = 0.24, p-val = 0.63). However, during the After trials, there was a negative relationship between the level of 17α,20β-P and both the number of contacts (F_(1,17)_ = 5.94, p-val = 0.02, Fig. 2) and the willingness to forage under risk (F_(1,17)_ = 8.02, p-val = 0.01, Fig. 2). Visual inspection of the data suggests that this pattern was particularly strong among females that had an opportunity to mate (Fig. 2). We did not detect a relationship between activity and 17α,20β-P (respectively, F_(1,17)_ = 0.21, p-val = 0.64).

**Figure 2.**
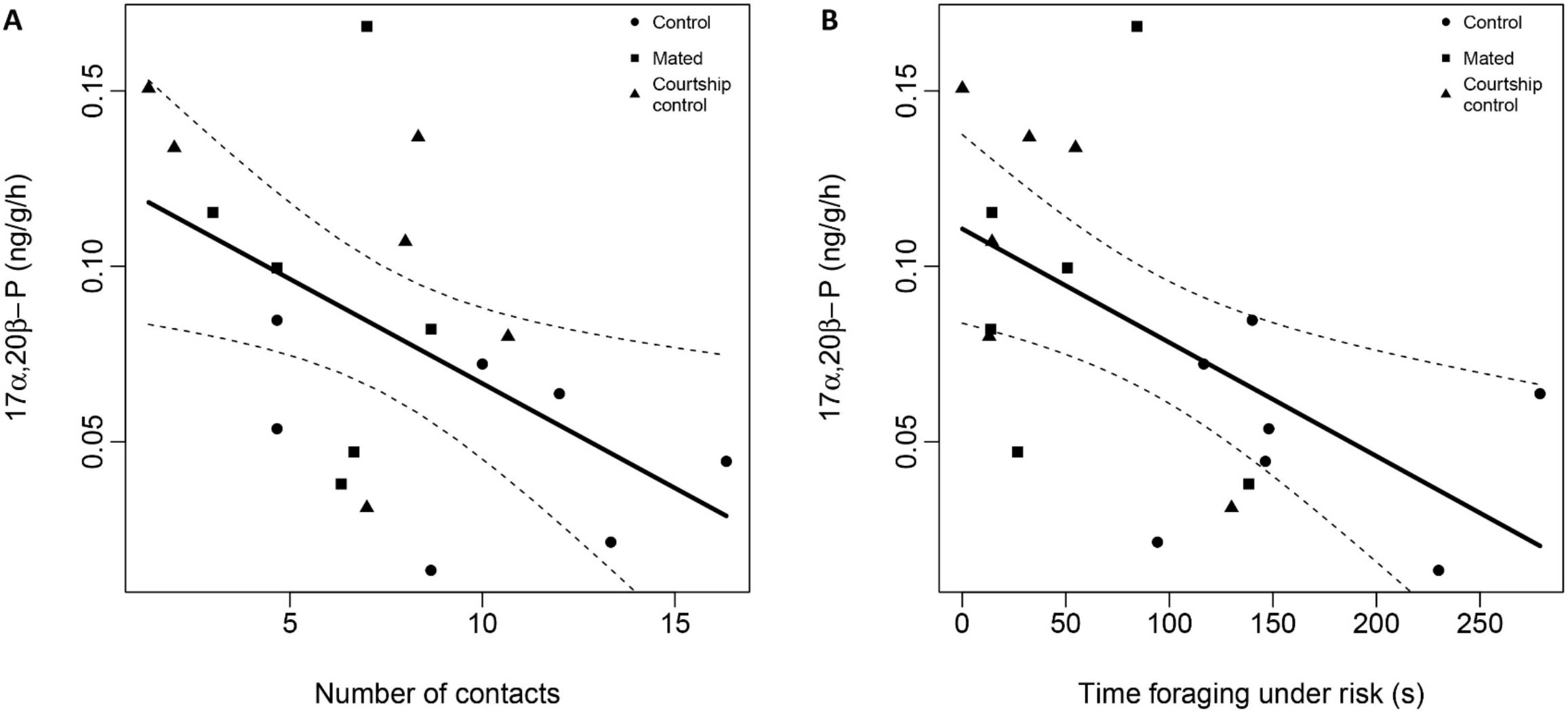
Relationship between progesterone and A) the social behavior (number of contacts), and B) the risk taking behavior (time foraging under risk) during the “After” trials. Each circle represents the average time an individual spent foraging under risk during the three “After” trials. The dashed line represents the 95% confidence interval.

Non-significant results about other steroids are in the electronic supplementary material.

## DISCUSSION

This study addresses important questions about personality change throughout the lifespan by analyzing the stability and change of personality traits in a controlled, randomized and longitudinal experiment. Specifically, we analyzed how repeatability, within and among individual variation in personality, and mean level changes in personality traits depend on a key life event: mating for the first time. We provide evidence that adult females became less social and less willing to take risks after a mating opportunity.

This conclusion is bolstered by the study’s strong experimental design. We confirmed that there are consistent individual differences in behavior by repeatedly measuring the same individuals in a battery of behavioral assays. The repeated measures design revealed that individuals given a mating opportunity retained their new behavioral type for the duration of the experiment. Moreover, it allowed us to compare within- and among-individual variance components, revealing greater within-individual variation in individuals given a mating opportunity compared to the control group. This is strong evidence that the mean-level differences between treatment groups reflects within-individual change, i.e. individuals changed their behavior after a mating opportunity. In addition, we confirmed that self-selection was not a problem: at the beginning of the experiment, females in the three treatment groups did not differ in behavior, but they diverged following the mating opportunity.

Interestingly, regardless of whether they physically mated or if they just had the social experience of courtship, females became less social and less willing to take risks after a mating opportunity. We do not know why some females given a mating opportunity mated and others did not. It is possible, for example, that mated females experienced less aggression from males during the mating opportunity, that mated females were more attractive to males, that mated females were less choosy, or less interested in mating generally. However, prior to entering the experiment, we did not detect any differences in behavior or body size or reproductive state (gravidity) between mated and courtship control females (electronic supplementary material), both types of females were given numerous opportunities opportunities to mate. There was not a systematic difference between the males that were offered to females that did versus did not mate, and we ensured that all females were gravid and ready to spawn when given a mating opportunity. Mate choice in sticklebacks is mutual [27]; therefore, we suspect that some females did not mate simply because they were not offered the right match. Despite the fact that mated vs courted females might have differed in attractiveness, choosiness or the way they interacted with males, both types of females became less social and less willing to take risks and experienced an increase in progesterone. These results strongly suggest that there is something about the experience of having an opportunity to mate with males that changes females’ personality, i.e. the social experience is sufficient to cause personality traits to change. That mating and experiencing courtship do not differ in their impact on female personality might stem from the fact that mating in this system is similarly costly to experiencing courtship. The similar effects on personality may be from the similar physiological impacts of the behaviors involved in each case; a different outcome might be expected when the act of mating for females is more costly than engaging in courtship only. If that is the case, then the impact of a mating opportunity on personality in this study may be a conservative example of the impact of mating when considering systems in which the females then go on to perform parental care.

The similarity between courtship control and mated females in this experiment suggest that social experience with potential mates – not just sexual experience – can influence personality traits in females. This result is consistent with what we know about the importance of social experience for behavior in both humans and nonhuman animals. For example, for young adults, people’s openness and their interaction with their social environment influences their chances of meeting a partner [28,29]. The nonhuman animal literature is rife with examples showing that previous courtship experience with male signals alters female mating decisions [13,30]: females are often more choosy about their mates after social experience with an attractive male and less choosy after social experience with an unattractive male, presumably because females change their preference functions as they update their estimate of the distribution of mate quality [31]. Indeed, there is evidence that female sticklebacks modify their mate preference in response to their estimate of the quality and availability of mates [32,33]. Therefore, it is possible that females used the mating opportunity in this experiment to update their assessment of mate availability, which went on to influence their social and risk taking behavior.

Females excreted more 17α,20β-P after they were given a mating opportunity, regardless of whether they mated, and individual variation in behavior was related to individual variation in 17α,20β-P. 17α,20β-P promotes the secretion of ovarian fluid in sticklebacks [34] and it is likely that it is the maturation inducing hormone in stickleback [24]. In other fishes, a sharp peak of 17α,20β-P occurs prior to spontaneous or induced ovulation [35,36]. Studies in goldfish and other fishes have shown that exposure to a potential mate can trigger ovulation, presumably via pheromones [37], and there is some evidence for chemical communication between males and females during courtship in stickleback [38]. Therefore, the higher levels of 17α,20β-P in females given a mating opportunity in this experiment could reflect their recent experience with a potential mate. Alternatively, because 17α,20β-P can act as a pheromone used during communication among females [39], higher levels of 17α,20β-P in both the mated and courtship control groups might reflect 17α,20β-P that was released by females that mated while they were co-housed with courtship control females. Interestingly, contrary to our prediction based on the coping styles literature [23], cortisol did not appear to be involved in mediating the effects of a mating opportunity on personality.

Regardless of the proximate physiological mechanisms involved, the finding that risk taking behavior and social behavior decreased following a mating opportunity is consistent with theory which posits that life-history tradeoffs - specifically between investment in current and future reproduction - can generate behavioral types [14]. Specifically, individuals with a greater expectation of future reproduction are expected to be shyer and less aggressive than individuals with a lower expectation of future reproduction [14]. In the context of this experiment, females that had an opportunity to mate may have interpreted this experience as information that potential mates are abundant, therefore these females may have a higher expectation for future reproduction, and thus became less social and less willing to take risks in order to protect their assets [15].

It is now a truism that early life experience is important for behavioral development in humans and nonhuman animals [4,40], but whether and why personality traits continue to change through adulthood is less understood. Experimental studies that manipulate and control particular life events in nonhuman animals have the potential to make important contributions in this area. Moreover, this topic deserves more attention in the animal personality literature because it has a number of ecological and evolutionary implications that have yet to be explored. For example, personality change in young wild individuals may have strong consequence for dispersal and/or the establishment of social groups, with potential consequences for social interactions among individuals within the group. More generally, these results highlight the importance of phenotypic plasticity over ontogeny for the generation and maintenance of personality variation within natural populations; changes in the timing of important life history events or other experiences could have consequences for the distribution of personality variation within natural populations.

## Ethics

All applicable international, national, and/or institutional guidelines for the care and use of animals were followed. All procedures performed in this study involving animals were in accordance with the ethical standards of the University of Illinois, Urbana Champaign (IACUC protocol #18080).

## Data accessibility

Data available from the Dryad Digital Repository at: Digital Repository at: https://doi.org/10.5061/dryad.wstqjq2h3 [41].

## Authors’ contributions

C.M. contributed to study design, carried out the experiment and the laboratory work, analyzed the data, and co-wrote the manuscript; A.M.B. contributed to study design and co-wrote the manuscript.

## Acknowledgements

The authors thank Dana Joulani, Katie Julkowski, Jason Keagy, Lucas Li, Ilva Mane, Ryan Paitz, Jake Ritthamel, and Rosie Zhang. This work was supported by a fellowship from the Fyssen foundation to CM, grants from the NIH (2R01GM082937-06A1) and NSF (IOS 1121980), and the University of Illinois at Urbana-Champaign.

## Supplemental material

**Figure S1.**
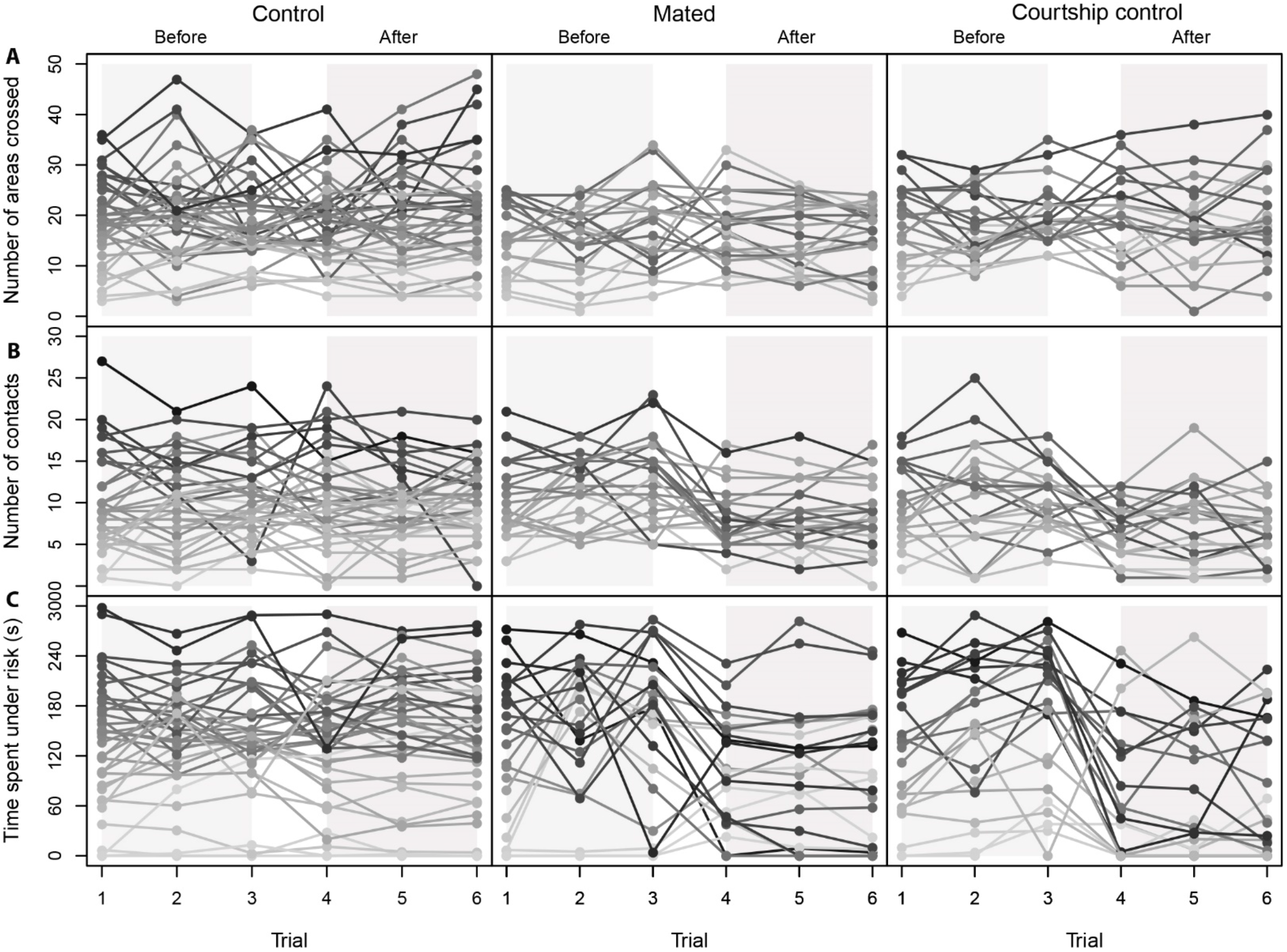
Behavioral reaction norms showing individual differences in behavior the three treatment groups. Top to bottom shows: A) number of areas crossed during the activity assay, B) number of contacts during the social behavior assay and C) time spent foraging under risk during the risk taking behavior assay. Each line represents the behavior of a different individual female across all six trials in shades of grey. Trials 1-3 represent behaviors measured during the “Before” trials, trials 4-6 represent behaviors measured during the “After” trials.

**Figure S2.**
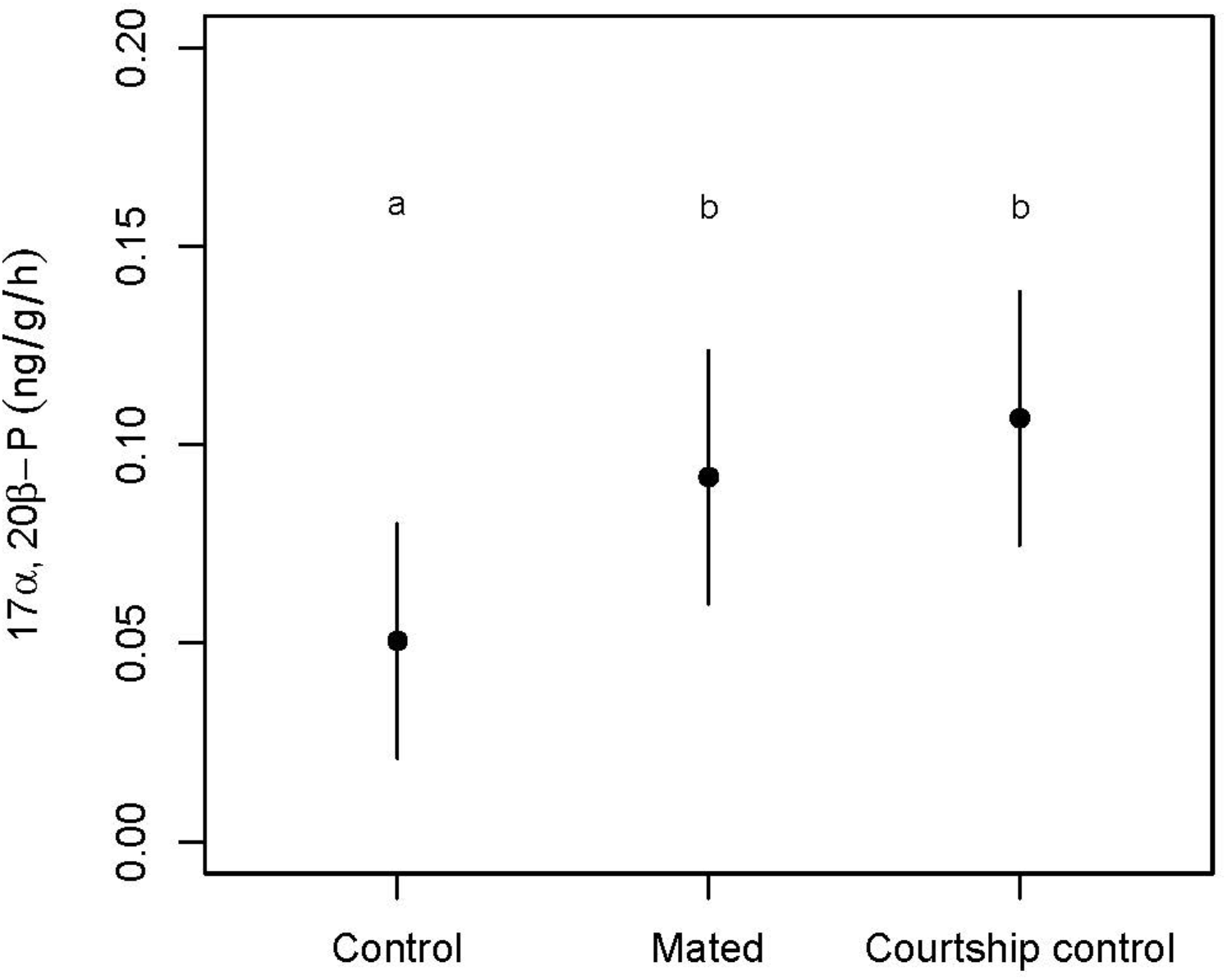
Differences in average excreted progesterone among treatment groups. Shown are the “After” progesterone levels. Bars represent the 95% confidence interval. Different letters indicate means that are significantly different (p-val < 0.05).

**Figure S3.**
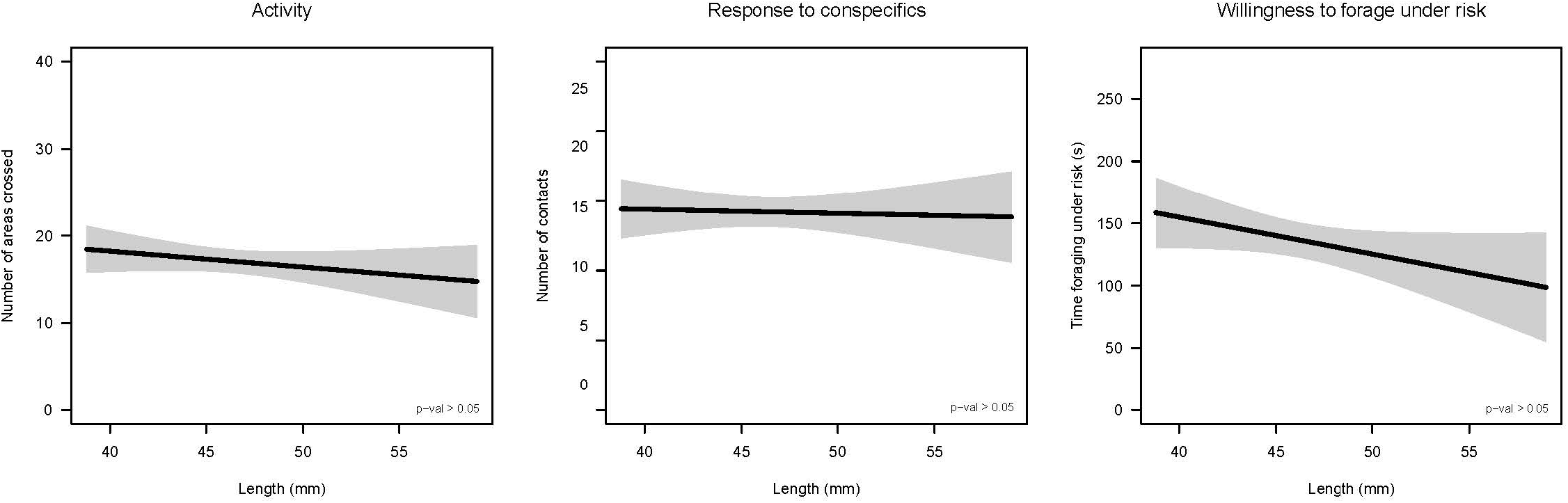
There was no relationship between body size (standard length) and behavior (A: activity, B: social behavior, C: risk taking behavior). The graph shows the predicted linear regression between length and each behavior, and the associated confidence interval (95%). Shown from left to right are the number of areas crossed during the activity assay, the number of contacts during the social behavior assay, and the time foraging under risk during the risk taking assay.

**Table S1.** Repeatability (R) and variance component (among- and within-individual) of behavioral traits during the “Before” trials. For both control females and females in the mating opportunity treatment we estimated the repeatability of activity (number of crossed areas), their social behavior (number of contacts), and their risk taking behavior (time foraging under risk) across the three “Before” trials. Models included individual as a random effect and adjusted models included trial as a co-variate. Numbers in brackets indicate 95% credibility intervals (82 individuals with 3 repetitions). Repeatability, among- and between-individual variation did not differ between the two groups prior to the experience of courtship and reproduction.

|  | Mating<br>opportunity | Control |
| --- | --- | --- |
| **Number of areas crossed** |  |  |
| Among | 26.17 | 44.72 |
| Within | 24.88 | 35.66 |
| R | 0.463 (0.28, 0.62) | 0.55 (0.34, 0.69) |
| **Number of contacts** |  |  |
| Among | 13.47 | 15.32 |
| Within | 9.52 | 7.70 |
| R | 0.58 (0.40, 0.72) | 0.66 (0.48, 0.76) |
| **Time foraging under risk** |  |  |
| Among | 4435.90 | 4425.90 |
| Within | 2858.50 | 757.10 |
| R | 0.59 (0.43, 0.81) | 0.85 (0.75, 0.91) |

**Table S2.**
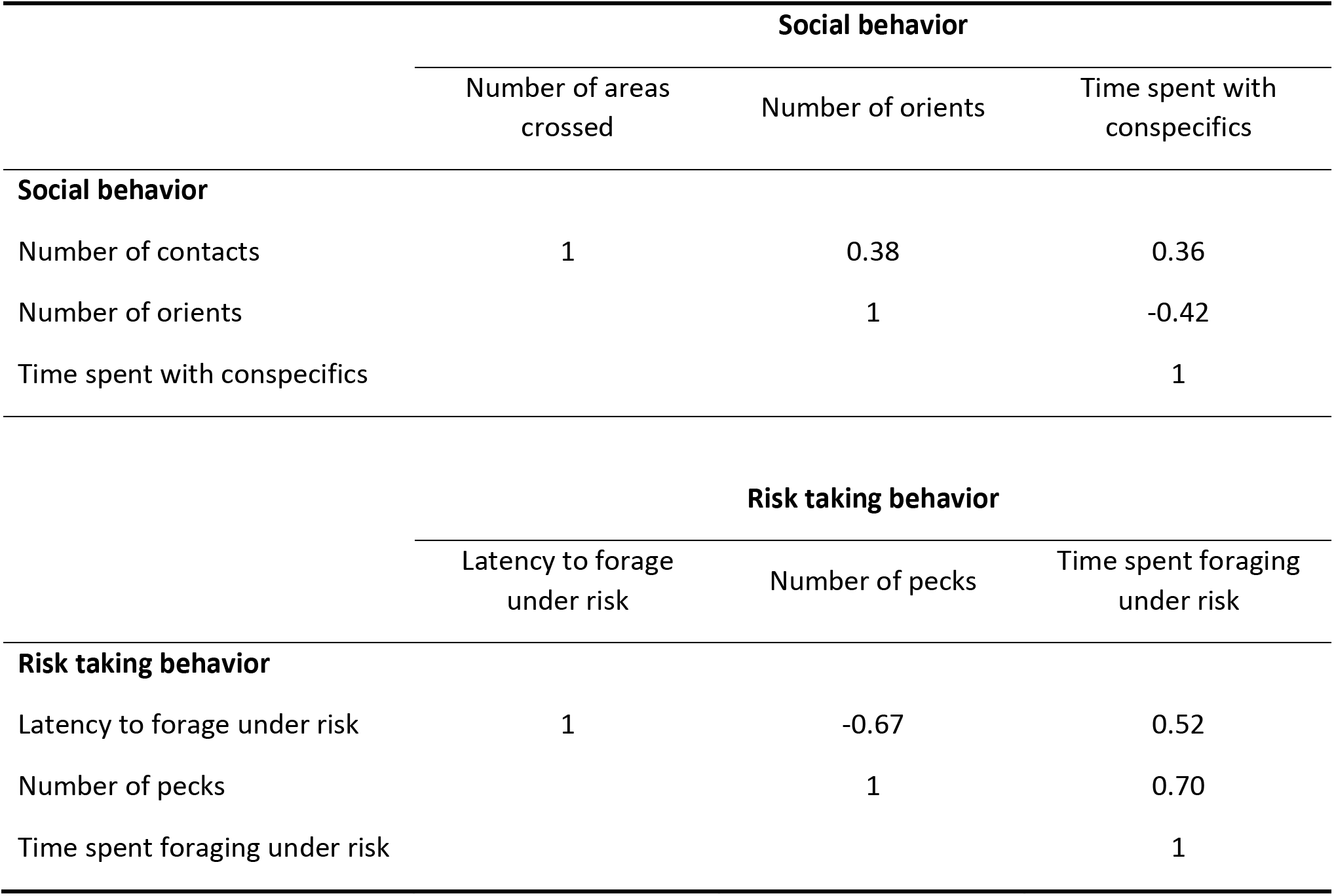
Matrix of the Spearman’s correlation coefficients among behaviors for the social behavior and the risk taking behavior assays; behaviors were averaged across the six trials (n=82 individuals).

|  | **Social behavior** |  |  |
| --- | --- | --- | --- |
|  | Number of areas crossed | Number of orients | Time spent with conspecifics |
| **Social behavior** |  |  |  |
| Number of contacts | 1 | 0.38 | 0.36 |
| Number of orients |  | 1 | -0.42 |
| Time spent with conspecifics |  |  | 1 |
| **Risk taking behavior** |  |  |  |
|  | Latency to forage under risk | Number of pecks | Time spent foraging under risk |
| **Risk taking behavior** |  |  |  |
| Latency to forage under risk | 1 | -0.67 | 0.52 |
| Number of pecks |  | 1 | 0.70 |
| Time spent foraging under risk |  |  | 1 |

