## Supplemental Material for "Personality traits change after an opportunity to mate"

### Electronic supplementary material

#### METHODS

##### Overview of the experiment

In order to track changes in personality as a function of mating, the behavior and excreted hormone levels of adult female threespine sticklebacks was recorded both before and after mating, and compared relative to females that did not have an opportunity to mate (control) and to females that had an opportunity to mate and therefore experienced courtship socially but did not mate (courtship control). Individual behavior was repeatedly recorded in three separate behavioral assays designed to measure activity, social behavior and risk taking behavior. The behavior of individuals in all three treatment groups was measured in each assay three times both before and after females had an opportunity to mate, for a total of six trials per assay per individual. Repeated samples of steroids excreted in water were collected and were measured using GC/MS.

##### Study population

Threespine sticklebacks were collected from Putah Creek, California, U.S.A. in April 2017 prior to the onset of the breeding season. Neither males nor females showed signs of reproductive maturity when they were collected; therefore, we assume that females were virgins at the time of the experiment. Fish were shipped to the University of Illinois at Urbana-Champaign (Champaign, IL, U.S.A.) and maintained in the laboratory in mixed-sex group tanks (108 L × 33 W × 24 H cm). Fish were housed in tanks (60.75 L × 30 W × 20 H cm, 20 fish/tank) for approximately three weeks before the experiment began.

One week before the start of the experiment, females were housed in 'home' tanks (35.5 L × 33 W × 25 H cm, 10 fish/tank) with a gravel bottom, plastic plants, and a shelter (opaque white PVC pot of 10 cm in diameter) and stayed in these home tanks for the duration of the experiment, except when individuals were transferred to separate 'observation' tanks (60.75 L × 30 W × 20 H cm) for behavior trials. The observation tanks were set up the same as the home tanks, i.e. gravel at the bottom, plastic plants, and a shelter (white PVC pot). All trials were conducted from mid-June to early October 2017 i.e. during the reproduction period of this population. None of the females in this experiment were infected with *Schistocephalus solidus*, a tapeworm known to influence risk-taking behavior [1,2].

Fish were maintained at 20°C on a summer photoperiod (16: 8 h light: dark cycle). A recirculating flow-through system consisting of a series of particulate, biological, and UV filters (Aquaneering, San Diego, USA) was used to clean the water. 10% of the water volume in the tanks was replaced each day. The fish were daily fed a mixed diet consisting of frozen bloodworm, frozen brine shrimp, and frozen Mysis shrimp ad lib each day. During behavioral assays, fish were fed after assays ended.

**Experimental design**

Individual females were randomly assigned to either the control group or to have a mating opportunity. Females given a mating opportunity were paired with a control female (matched for size) who was always measured at the same time as her partner. This experimental design allowed us to control for variation among females in time to become gravid and to reproduce, as well as for self-selection, i.e. if some behavioral types of females are more willing to mate or more attractive than others. Females in the control group (n = 37) were not exposed to a male or given an opportunity to mate. Some (n = 23) of the females given a mating opportunity mated, while others (n = 22) did not and served as “courtship controls”, because like the mated females, they had the social experience of being courted, but unlike the mated females they did not physically mate. We did not detect any latent differences between females that mated and courtship control females that would lead to nonrandom representation of behavioral types between the mated and courtship control groups, e.g. mated and courtship control females did not differ in size ($t_{43.95}$= 0.16, P = 0.87) or behavior (see Results).

Males were selected for this experiment according to throat and eye color. All of the females that were introduced in a tank with a male were gravid and ready to spawn. If a female did not mate with a male, she was given another opportunity to mate with a different male. We usually had several females and males ready to mate at the same time so that we first randomly picked the first male-female pairs and then, if they did not mate, we crossed pairs. Therefore, both courtship control and mated females encountered the same males on average and they were given the same opportunity to mate.

The control females were in isolation while their paired female was given a mating opportunity therefore it is conceivable that the differences between control females and females given a mating opportunity reflects the opportunity for a social interaction in general, not the experience of interacting with a male per se. However, this is highly unlikely because all of the females in this experiment had an opportunity to interact with other females at multiple points during the experiment. That is, females were in social groups with other females for almost the entire experiment; the only time that females were apart from conspecifics was when they were going through the behavioral assays and when females were given an opportunity to mate. Therefore, there was an opportunity for social interactions among females to influence the subsequent behavior of females in all three conditions. Given equal opportunity for social interactions among females to influence behavior in all of the different conditions, it is highly unlikely that the differences between females given a mating opportunity and control females reflects the opportunity for social interactions per se.

The behavior of females was measured in the behavioral assays (described below) on six occasions, thrice in the “Before” trials and thrice in the “After” trials. After the third and last day of the “Before” trials, we transferred females to separate home tanks with the same characteristics as the initial home tank to wait for them to become gravid. Gravidity was determined by the presence and position of the abdominal distension and the fact that we were able to see the eggs through the body wall. As fish were paired, we also transferred the paired-control female at the same time in a similar new home tank, where they remained while their paired partner had an opportunity to mate once she became gravid. Females given a mating opportunity were placed in a tank with a male who had built a nest (60.75 L × 30 W × 20 H cm). If the pair did not mate within 30 minutes, the female was removed from the male’s tank and was allowed to recover for at least three hours before being introduced to a different male’s tank for another mating attempt. The number of mating attempts until successful reproduction varied between 1 to 8, mean ± SE = 2.9 ± 0.29. We did not detect an effect of the number of mating opportunities on behavior (activity: $F_{(1,268)}$ = 0.048, p-val = 0.83; social behavior: $F_{(1,268)}$ = 0.031, p-val = 0.86; risk-taking behavior: $F_{(1,268)}$ = 3.055, p-val = 0.062). Therefore courtship control females were given at least one opportunity to mate but for whatever reason (as discussed above), they did not. Females were never prevented from mating. Mated females were weighed before and after mating in order to estimate clutch size. Females in all three treatment groups often became gravid over the course of the experiment. In order to control for the effects of gravidity on behavior [3–5], we waited until either the female spawned with a male or naturally lost her eggs (i.e. eggs were found in the female’s tank a couple of hours after her last mating opportunity, the female did not show morphological signs of gravidity and the female weighed less than before) before running her through the behavioral assays. Therefore, females were never gravid during the behavioral assays.

**Behavioral assays**

*Activity*

The focal fish was placed in an opaque shelter (white pot with a removal cork) at one end of the tank. After one minute, the observer gently removed the cork of the shelter, and recorded the latency to emerge from the shelter (“latency to emerge”). Then, the observer recorded the number of times the fish froze (i.e. the fish maintains the same position without moving, “number of freezes”), and the total number of times the fish crossed into a different area (“areas crossed”, 4 areas of about 15 cm each) for three minutes. Short latencies to emerge, few freezes and large numbers of areas crossed were interpreted as relatively high levels of activity and exploratory behavior. Initial analyses showed that the latency to emerge and the number of freezes were not discriminant enough, so we chose to focus on the number of areas crossed in the final analyses.

*Social behavior*

30 minutes after the behavior of the focal fish was measured in the activity and exploratory behavior assay, three females from the same population were placed into a flask at the opposite side of the refuge and the following behaviors of the focal fish were recorded for 5 minutes: the number of times the focal fish approached the conspecifics within one body length (“contacts”), the number of times the focal fish oriented to the conspecifics while more than one body length away from the conspecifics (“number of orients”), and the total amount of time (in seconds) the focal fish spent within one body length of the conspecifics oriented to conspecifics or not (“time with conspecifics”). We interpret high rates of biting as aggressive behavior, and high rates of contacts, orienting and time with the conspecifics as high social behavior. Initial analyses showed that the behavioral variables were correlated with one another (electronic supplementary material, Table S2), and that number of contacts was more amenable to linear analyses therefore we chose to focus on the number of contacts in the final analyses.

*Risk taking behavior*

30 minutes after the behavior of the focal fish was measured in the social behavior assay, a model great egret (*Casmerodius albus*) head was placed over the observation tank. Then, we added live bloodworms directly under the egret. When the focal fish approached the worms within one

body length, we released the egret twice in quick succession, and we then fixed the egret so that it remained above the water. Following this simulated attack, the following behaviors of the focal fish were recorded for five minutes: the latency (in seconds) to the first bite at the worms after the egret strike (“latency to forage under risk”), the number of pecks at the worms (“number of pecks”), and the total time (in seconds) the fish spent foraging within one body length of the worms (“time foraging under risk”). We interpret a short latency to resume foraging and high rates of foraging following the simulated egret attack as relatively ‘bold’ behavior. Following this trial, the focal fish was removed from the observation tank and returned to its home tank. Initial analyses suggested that these behavioral variables were correlated with one another (electronic supplementary material, Table S2), and that time foraging under risk was more amenable to linear analyses, therefore we chose to focus on time spent foraging under risk in the final analyses.

The behavior of each female in each assay was assessed three times before the mating opportunity (Before trials), and three times after the mating opportunity (After trials), with 24 h between trials. Females started the After behavioral assays 24 hours after the mated female reproduced or after the courtship control female lost her eggs in her home tank.

The interval between the “before” and “after” trials varied between 7 to 45 days (mean ± SE = 18.64 ± 1.34). The number of days that elapsed between the “before” and “after” trials differed among females because we waited for females to naturally become gravid. We elected to follow the female’s timeline rather setting an arbitrary interval between the before and after trials in order to be able to pinpoint the cause of changes in personality, i.e. due to the social experience of being courted/mated.

All females were measured for length and weight on the last day of the After trials.

**Measuring excreted steroids**

After the third trial of both the Before and After trials, we placed the focal fish in a 500 ml long-necked glass flask filled with 100 ml of water. We then placed the flask in a covered bucket (to minimize stress) for 30 minutes. Then, we transferred 50 ml of the water into a 50 ml sterile polypropylene conical tube. Water samples were collected between 11.00 am and 04.00 pm

Central Standard Time. Water samples were frozen at -20°C until extraction. Freeze-storage of water samples does not influence steroid concentrations [6].

Steroids were extracted from the water samples by pulling water through C18 Sep-Pak cartridges (Waters Ltd.) that had been primed with 5 mL methanol followed by 5 mL distilled water. After the sample had dripped through at a rate of approximately 2 mL/min, the cartridge was washed with 5 mL of distilled water, and the steroids were eluted from the columns into 13*100 mm borosilicate vials via 5 ml of diethyl ether. The ether was dried by evaporation overnight. The dried hormones were then frozen at -80°C to preserve them following the advice of the Metabolomics Center (Roy J. Carver Biotechnology Center, University of Illinois at Urbana-Champaign).

Samples were analyzed via mass spectrometry with the 5500 QTRAP LC/MS/MS system (AB Sciex, Foster City, CA). The 1200 series HPLC system (Agilent Technologies, Santa Clara, CA) includes a degasser, an autosampler, and a binary pump. The LC separation was performed on a Phenomenex C6 Phenyl column (2.0*100 mm, 3 µm) with mobile phase A (0.1% formic acid in water) and mobile phase B (0.1% formic acid in acetonitrile). The autosampler was set at 5°C. The injection volume was 5 µL. Mass spectra was acquired under positive electrospray ionization (ESI) with the ion spray voltage of 5500 V. The source temperature was 500°C. The curtain gas, ion source gas 1, and ion source gas 2 were 36 psi, 50 psi, and 65 psi, respectively. Multiple reaction monitoring (MRM) was used to measure hormones with the Q1–Q3 transition of 363.1–121.0 (m/z). In order to control for differences in body size, hormone release rates were calculated as the amount of released hormone per gram of body weight per hour (ng/g/hr). We focus on 17α,20β-P (17α,20β-dihydroxy-4-pregnen-3-one referred as 17α,20β-P in the text, n = 19 individuals with one measure Before and one measure After), estradiol (n = 65) and cortisol (n = 49). 17α,20β-P stimulates the production of ovarian fluid [7] and is likely to be the maturation inducing hormone in stickleback [8], estradiol increases after spawning, stimulating oogenesis and then drops when the eggs are ready to be ovulated [9] and cortisol is released in response to stress and often studied in the context of personality trait variation [10,11].

#### Statistical analyses

##### *Repeatability of behavior and hormones*

To confirm that there were consistent individual differences in activity, social behavior and risk taking behavior, we estimated the repeatability of behavior during the Before trials. We measured repeatability for the three treatment groups separately to confirm that all groups showed similar patterns prior to the mating opportunity. To test whether mating influences rank-order stability (i.e. among-individual variation), we estimated the repeatability of behavior and hormone titres across the six Before and After trials for the treatment groups separately.

Repeatability was estimated as the ratio of between-individual variance to total variance with linear mixed-effects models (with individual identity as a random factor) in R v.3.4.4 (http://www.r-project.org) [12]. To determine whether repeatability estimates were significantly different from zero, we estimated the upper and lower 95% confidence intervals and visually inspected whether they were pressed against zero. We also determined whether repeatability estimates were significantly different among treatment groups by looking at whether the 95% confidence intervals overlapped.

##### *How do personality traits and hormones change following mating?*

To determine whether there was mean-level personality trait change after mating, we used linear mixed-effects models (LMMs) with the lmer function in the R package lme4 v.1.1-17 [13]. Models examining mean-level change over time included the following fixed effects: treatment (three levels: control, courtship control, and mated), Before/After (two levels: before vs after mating), trial nested within Before/After, the number of days between the first and sixth trial (days in the experiment) and the interaction between treatment x Before/After. All models included individual as a random effect to account for the six measurements of the same subjects. Body size was not included in the final analyses because we did not detect a difference in size among treatments and we did not detect correlations between size and any of the behavioral variables (see electronic supplementary material, Figure S3).

For the hormonal release rate, we split the all data set by the period (Before/After) as we did not have enough statistical power to test for the interactions between factors such as treatment and period. Therefore, we ran simple linear regressions with individual as a random effect. Secondly, following the same rule, we performed linear models to investigate a potential link between hormones and personality traits. To avoid pseudoreplication, we first averaged each behavior across the three trials within either the before or after trials.

#### RESULTS

##### Effects of a mating opportunity on behavior

*Effects of other predictor variables on behaviors*

There was no evidence that behavior systematically changed over time, i.e. no evidence for habituation across the repeated trials, as there was no effect of trial on behavior (number of areas crossed: $F_{(2,489)} = 0.93$, p-val = 0.40; number of contacts: $F_{(2,489)} = 0.44$, p-val = 0.64; willingness to forage under risk: $F_{(2,489)} = 0.77$, p-val = 0.46). Importantly, the number of days that elapsed between the before and after trials had no effect on behavior (number of areas crossed: $F_{(1,490)} = 1.23$, p-val = 0.27; number of contacts: $F_{(1,490)} = 2.02$, p-val = 0.15; willingness to forage under risk: $F_{(1,490)} = 1.67$, p-val = 0.20).

##### Effects of a mating opportunity on hormones

*Effects of predictor variables on cortisol and estradiol*

Neither time nor treatment affected excreted cortisol or excreted estradiol (Before/After (cortisol: $t_{96} = 0.87$, P = 0.38; estradiol: $t_{128} = -0.43$, P = 0.66, treatment (cortisol: $F_{(2,95)} = 0.04$, p-val = 0.96; estradiol: $F_{(2,127)} = 1.03$, p-val = 0.36).

##### Relationship between hormones and behavior after a mating opportunity

*Relationship between cortisol or estradiol and the three behaviors*

We did not detect a relationship between cortisol or estradiol and any of the behaviors (activity and exploration: cortisol: $F_{(1,47)}$ = 0.63, p-val = 0.43; estradiol: $F_{(1,63)}$ = 0.30, p-val = 0.59; social behavior: cortisol: $F_{(1,47)}$ = 1.16, p-val = 0.29; estradiol: $F_{(1,63)}$ = 1.09, p-val = 0.30; risk taking behavior: cortisol: $F_{(1,47)}$ = 1.08, p-val = 0.30; estradiol: $F_{(1,63)}$ = 0.11, p-val = 0.75).
